# Cell-based production of Fc-GlcNAc and Fc-alpha-2,6 sialyl glycan enriched antibody with improved effector functions through glycosylation pathway engineering

**DOI:** 10.1101/2023.12.18.572280

**Authors:** Han-Wen Huang, Vidya S. Shivatare, Tzu-Hao Tseng, Chi-Huey Wong

**Affiliations:** Department of Chemistry, The Scripps Research Institute, La Jolla, California 92037, United States

## Abstract

Glycosylation of antibody plays an important role in Fc-mediated killing of tumor cells and virus-infected cells through effector functions such as antibody-dependent cellular cytotoxicity (ADCC), antibody dependent cell-mediated phagocytosis (ADCP) and vaccinal effect. Previous studies showed that therapeutical humanized antibodies with α2-6 sialyl complex type (SCT) glycan attached to Fc-Asn297 exhibited optimal binding to the Fc receptors on effector cells associated with ADCC, ADCP and vaccinal effect. However, the production of antibodies with homogeneous Fc-SCT needs multiple *in vitro* enzymatic and purification steps. In this study, we report two different approaches to shorten the processes to produce SCT-enriched antibodies. First, we expressed a bacterial endoglycosidase in GNT1-KO EXPI293 cells to trim all *N*-glycans to mono-GlcNAc glycoforms for *in vitro* transglycosylation to generate homogeneous SCT antibody. Second, we engineered the glycosylation pathway of HEK293 cells through knockout of the undesired glycosyltransferases and expression of the desired glycosyltransferases to produce SCT enriched antibodies with similar binding affinity to Fc receptors and ADCC activity to homogenous SCT antibody.

## Introduction

Antibody plays an important role in humoral immune response and has diverse plasticity and biological functions. The prototype of antibody is a major component of B cell receptor (BCR) on naive B cells. When the BCRs are activated through pattern recognition receptors (PRRs), the alternative splicing of mRNA helps produce antibodies as secreted forms of IgM or IgD. With the assistance of dendritic cells and T helper cells, the activated B cells could further undergo clonal expansion, somatic hypermutation, and class switch and are differentiated into plasma cells to produce secreted IgG, IgA, or IgE antibodies with high antigen specificity. Once a target cell or pathogen is opsonized by an antibody, it may trigger different downstream immune functions, including phagocytosis, cellular cytotoxicity, vaccinal effect, complement activation etc. These immune cell-based responses required the binding of antibody Fc domain to specific Fc receptors on immune cells. Each subtype or glycoform of antibody can be recognized by several Fc receptors rendering activating or inhibitory signaling to generate different effector functions. For example, the binding of IgG antibody to the FcγIIIA receptor on NK cells will activate NK cells to release cytokines to kill target cells, a process called antibody-dependent cellular cytotoxicity (ADCC)^1-3^ and binding to the FcγIIA receptor on macrophages will activate macrophage to kill the target cell through antibody-dependent cellmediated phagocytosis (ADCP). In addition, binding of antibody-antigen complex to the FcγIIA receptor on dendritic cells can facilitate dendritic cell maturation to induce memory CD8 T-cell response ^4^.

Glycosylation is a common post- or co-translational modification found on most cell surface proteins. During *N*-glycoprotein synthesis, a high mannose glycan is first synthesized in the ER and then transferred to the growing polypeptide chain followed by chaperone-mediated protein folding and quality control to form glycoproteins. The high-mannose glycoproteins are trimmed by mannosidases and translocated to the Golgi apparatus for addition of more monosaccharides sequentially by different glycosyltransferases before moving to the cell surface. Unlike protein and DNA, glycosylation is temple independent, and the sequence of glycans is determined by the repertoire of glycotransferases expressed in different cell types. Moreover, the cellular environment and protein accessibility also affect the activity and specificity of glycosyltransferases. Thus, the glycans generated by glycosylation usually exhibit more diversity than other modifications^5-7^.

Antibodies of each subtype are also glycosylated at different places within the constant region of heavy chain (HC) and these glycans regulate antibody functions^1^. The preBCR assembly is important for B cell development and is critically regulated by the *N*-glycan at N46 on μHC^8^. The *N*-glycan at N402 on μHC has been linked to antibody oligomerization and complement activation^9-10^. Besides, IgG *N*-glycosylation at N297 on γHC plays a critical role in complement activation and Fcγ receptor activation leading to various effector functions^11-13^. Therefore, it is important to identify optimized glycosylation to improve the efficacy and safety of therapeutic antibodies.

Generally, the production of recombinant antibody is often based on mammalian cell lines, such as NS0, 293T and CHO cells, to produce a mixture of glycoforms with core-fucosylated bi-antennary complex type glycans^14^ which are not optimal for binding to the FcγIIIA receptor, because of the inhibitory function of core-fucosylation^11^ or the off-target delivery caused by terminal galactosylation^15^. Therefore, efforts have been directed toward modification of the Fc-glycan on antibody to improve specific receptor binding and the corresponding effector function. One strategy used was to block the fucosylation pathway in cells. For example, using the cell line deficient in GDP-fucose synthesis could produce antibodies without core-fucosylation, leading to an increase in binding to FcγIIIA receptor thereby enhancing the ADCC activity^11, 16^. The emergence of precise gene editing tools such as Zinc Finger Nuclease (ZFN) or Clustered Regularly Interspaced Short Palindromic Repeats (CRISPR)/ CRISPR-associated protein 9 (Cas9) has facilitated the progress of this strategy^17^ and studies on glycosylation pathway engineering in cells to produce therapeutic proteins with improved efficacy have been reported^18-19^.

Another strategy to alter protein glycosylation is through *in vitro* glycoengineering by enzymes or chemical ligation. A common approach was endoglycosidase-mediated transglycosylation^7^. Usually, the antibody with high mannose type glycan derived from insect cells^20^, yeast^21^ or GNT1 knockout (KO) cells was treated with endoglycosidase H or S2 to generate mono GlcNAc- or GlcNAc-Fuc bearing glycoforms. The antibody with either of these two glycoforms is a good acceptor substrate for mutated endoglycosidase (glycosynthase)-mediated transglycosylation^7^ using a homogenous glycan oxazoline as donor substrate to generate a new glycan chain. So far, many glycans have been transferred to antibodies to study the functional impact on receptor binding and effector functions. The fully galactosylated complex type glycan without core-fucose revealed that the α2-6 linked SCT provides enhanced binding to FcγRIIIA and FcγRIIA receptors which are associated ADCC, ADCP and vaccinal effect^23-24^. Although this strategy could produce antibodies as homogenous glycoforms, the process requires multiple reaction and purification steps and the specificity of mutant endoglycosidases is limited. In this report, we used humanized antibody chMC183-70 that targets specific stage embryonic antigen 4 (SSEA4) on various cancer cells as a model to produce and evaluate the Fc-SCT enriched antibody in Fc receptor binding and activation of cellular cytotoxicity.

## Result and Discussion

### Development of Fc-GlcNAc antibody producing cells by CRISPR-Cas9

Antibodies are generally produced from mammalian cell lines which contain core-fucosylated complex type glycans^14^. To generate the Fc-GlcNAc antibody for *in vitro* transglycosylation, antibody glycans need to be trimmed by endoglycosidase and fucosidase to leave only the innermost GlcNAc as an acceptor^7^. We would like to shorten these steps by CRISPR-Cas9 to generate Fc-GlcNAc antibody with commercially available GNT1 KO EXPI293F cells. The *N*-glycan expressed by this cell line was high mannose type which was contributes significantly to improve antibody binding to the FcγIIIA receptor compared to the wild-type antibody^22-23^. However, it also facilitates the antibody clearance by liver cells through asialoglycoprotein receptor binding. Further studies preferentially cleaved by Endo H^25^ to generate Fc-GlcNAc antibody. Thus, we aimed to knock in Endo H in this cell line by CRISPR-Cas9. However, after transfection of CRISPR-Cas9 and donor plasmids, cells started to die with apoptosis and resulted in reduced cell survival. Fortunately, the survival rate started to rise after several days and returned to normal level again (Fig. S1). We then wondered whether these surviving cells still expressed Endo H to cleave the mannose type *N*-glycans on antibody. The antibodies expressed by the cells were analyzed by electrophoresis and the result showed that the antibody heavy chain (AbHC) expressed by these cells had faster mobility than AbHC from its parental cells (GNT1 KO) and most AbHCs were downshifted (Fig. 2A). This antibody was also subjected to transglycosylation assay and the efficiency was good (Fig. 2C). We then analyzed the glycans on this antibody by intact protein mass analysis, and the results showed that most antibodies contain only GlcNAc on each HC (Fig. 2B and Fig. S2). These results indicate that CRISPR-Cas9-mediated Endo H KI in GNT1 KO cells can serve as a platform to produce Fc-GlcNAc antibody for downstream application.

**Figure 1.**
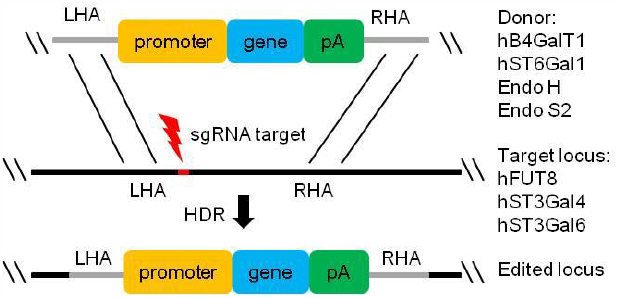
Illustration of CRISPR/Cas9 KO/KI strategies for glycosylation pathway engineering.

**Figure 2.**
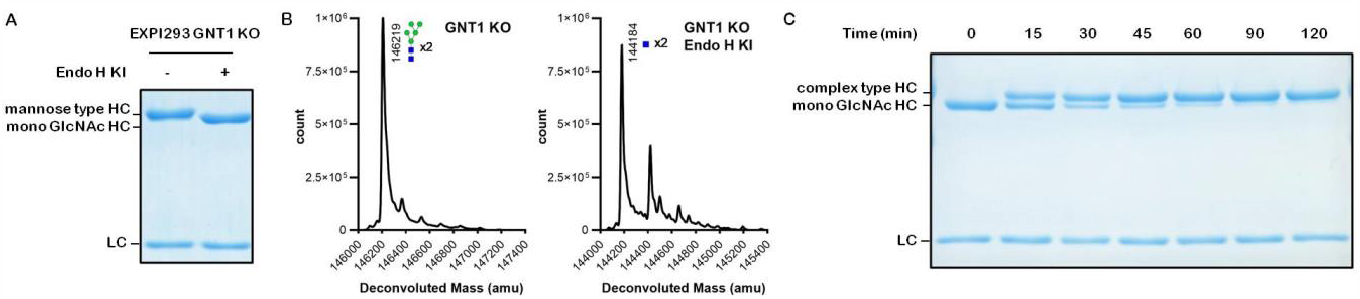
Expression of Endo H in GNT1-knock out cells results in cleavage of *N*-glycans on antibody. (A) The chMC18370 antibody expressed from engineered and its parental GNT1-KO EXPI293 cells were subjected to protein electrophoresis and the difference in molecular weight by band shifting was observed. (B) The chMC81370 antibody expressed by engineered and its parental GNT1-KO EXPI293 cells were subjected to IPM analysis to examine the molecular weight and glycan changes. (C) The chMC81370 antibody expressed by engineered GNT1-KO EXPI293 cells was subjected to transglycosylation assay and the efficiency of transglycosylation was assessed by protein electrophoresis.

In addition to Endo H, we also tried to express endoglycosidase S2 (Endo S2) from *Streptococcus pyogenes* in the cells because this enzyme can hydrolyze both high mannose and complex type *N*-glycans to Fc-GlcNAc^26^. Therefore, the donor vector and CRISPR-Cas9 plasmids were delivered to GnT1 KO cells and the cell survival rate was monitored. Interestingly, we did not see any significant cell death in the GNT1-KO cells after plasmid transfection (Fig. S1). Then the antibody glycans generated by these cells were examined as previously described.

The protein gel showed an obvious downshifted band of AbHC compared with its parental cells (GNT1 KO) (Fig. 3A). Besides, the glycans also can be added onto AbHC through enzymatic transglycosylation (Fig. 3C) and intact protein mass analysis showed a clear signal of Fc-GlcNAc antibody (Fig. 3B.). The glycoform analysis revealed a major Fc-GlcNAc antibody population as well (Fig. S2). Therefore, based on these results, it is concluded that endoglycosidase S2 can be introduced to cells by CRISPR-Cas9 to process the *N*-glycans to Fc-GlcNAc antibody for *in vitro* transglycosylation. In addition, this cell-based method can be used for other glycoproteins, such as influenza hemagglutinin and SARS-CoV-2 spike protein to generate the mono-GlcNAc decorated glycoforms as vaccines to elicit broadly protective immune responses^27-28^.

**Figure 3.**
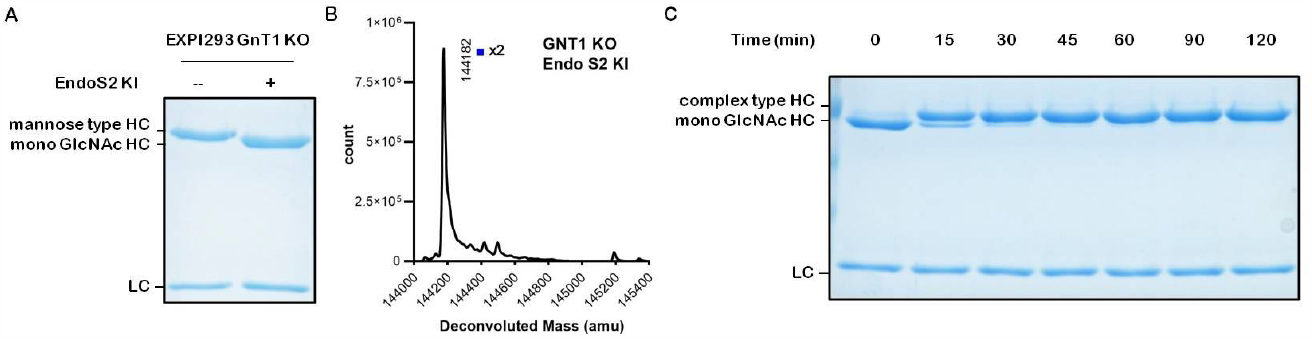
Expression of Endo S2 in GNT1-knock out cells results in cleavage of antibody *N*-glycans to GlcNAc. (A) The chMC18370 antibody glycoforms expressed by engineered and parental GNT1-KO EXPI293 cells were subjected to electrophoresis and a band shifting was observed. (B) The chMC81370 antibody expressed by engineered GNT1-KO EXPI293 cells was subjected to IPM analysis to examine the molecular weight and glycan changes. (C) The chMC81370 antibody expressed by engineered GNT1-KO EXPI293 cells was subjected to transglycosylation and the efficiency was evaluated by protein electrophoresis.

### Glycosylation pathway engineering in cells to produce Fc-SCT enriched antibody by CRISPR-Cas9

The second approach to generate antibody with Fc-SCT glycoform was to directly edit the expression of glycosyltransferases in cells. As previously described, the antibody produced from commonly used cell lines usually contained bi-antennary complex type *N-* glycans with core-fucose and terminated mainly with galactose^14, 18^-^19, 29^. Therefore, in order to produce the antibody with Fc-SCT, we decided to increase the level of galactosylation and sialylation on the antibody^19, 29^. In addition, the core-fucosylation, which depleted ADCC^11^, was eliminated by knocking out FUT8^30^ (Fig. 1). Furthermore, the terminal sialylation with α2-6 linkage, not the α2-3 linkage, was introduced as it was found to enhance receptor binding^31^ and improve ADCC and vaccinal effect^23-24, 32^. Although the mRNA of human α2-3 sialyltransferases could be detected in 293T and CHO cells^18, 33^, we did not observe any obvious sialylation on the antibody glycan from these cell lines^18-19, 29^ (Fig. 4, upper). Therefore, we were not concerned about the antibody being glycosylated by the inhibitory α2-3 sialylation. We therefore used CRISPR-Cas9 to knock out hFUT8 and knock in hB4GalT1 and hST6Gal1 in 293T cells (Fig. 1). The antibody expressed by this engineered cell line was purified and subjected to intact protein mass analysis of the antibody and the attached glycans. We observed a significant increase of galactosylated and sialylated antibody glycoforms (Fig. 4, lower) compared to the antibody generated by WT cells (Fig. 4, up). Glycoform analysis also revealed a significant increase in antibody galactosylation and sialylation, with lack of core-fucosylation (Fig. S3).

**Figure 4.**
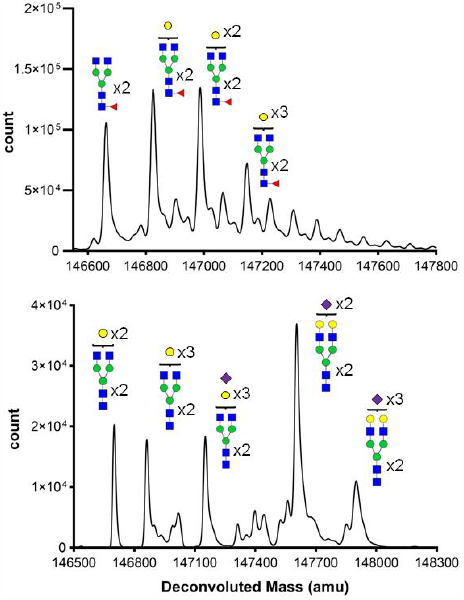
The glycoform composition of chMC81370 antibody. The *N*-glycans on antibody expressed by WT (upper) and glycosylation pathway engineered 293T cells (lower) were purified by protein A and analyzed by intact protein mass analysis.

### The sialylation on antibody is α2-6 linkage

To confirm the sialylation on the antibody was mediated by hST6Gal1, the antibody was treated with two different sialidases, *Streptococcus pneumonia* α2-3 neuraminidase and *Clostridium perfringens* neuraminidase overnight respectively. The intact protein analysis did not show any obvious difference in the distribution of asialylated and sialylated antibody between the untreated and α2-3 neuraminidase-treated group (Fig. 5 upper and middle), indicating that the sialylation on antibody was not due to the endogenous α2-3 sialyltransferases. On the other hand, using neuraminidase which could remove all linkages of sialylation largely eliminated the terminal sialic acids. In addition, the antibody with increased galactosylation generated from cells with knock in of hB4GalT1 only, the terminal sialylation was still not observed (Fig. S4). These results confirmed that sialylation on the antibody was introduced by hST6Gal1 and the increase of hB4Gal1 expression indeed elevated the galactosylation of antibody (Fig. 4). To further explore the capacity of sialylation or galactosylation in cells, we transiently overexpressed hST6Gal1 or hB4GalT1 in the engineered cells followed by antibody production to see if the galactosylated or sialylated antibody was increased. However, in intact protein mass analysis, we did not observe further elevation of any galactosylated or sialylated antibody after transient overexpression of hST6Gal1 or hB4GalT1 (Fig. S5). Another study was also attempted to increase the sialylation by elevating the glycosyltransferase expression in CHO cells but it still showed incomplete sialylated glycans^29^. The sialyltransferase hST6Gal1 is known as the only human glycosyltransferase which mediates α2-6 sialylation and preferentially use the α1-3 branch arm of the biantennary glycan as substrates^34^. The *in vitro* studies show glycoengineering on antibody glycans by hST6Gal1 takes a long time and difficult to yield fully sialylated antibody glycans^31, 34^. Furthermore, this also could explain why only the antibody with at least one fully galactosylated glycan was sialylated. If the galactose on G1 glycan was not on the α1-3 branch arm, the hST6Gal1 may not sialylate it. Therefore, the expression level of glycosyltransferases may not be the bottle neck in this situation rather the substrate preference and enzyme activity are the major factors^5-7^. Using α2-6 sialylatranferases from other organisms may have better activity than hST6Gal1.^35^

### Protein glycosylation varies with expression time

In order to understand if the amount and time of antibody overexpression in cells affected glycosylation, we extended the culture time to 5 days and then collected the medium at day 3 and day 5 to examine the antibody glycans by intact protein mass analysis. We unexpectedly found that the glycan pattern was somewhat different between these two time points (Fig. S6A). In the IPM analysis, a relatively strong sialylation was observed in the first collection (first three days) as we saw before (Fig. S6A, D0-3) and then sialylation was decreased in the second collection (last two days), accompanied by an increase of terminal gal-actosylation (Fig. S6A, D4-5), suggesting that the proteins involved in glycosylation may not be sufficient to effectively produce Fc-SCT antibody at later stage. Because the hST6Gal1 and hB4GalT1 involved in the FC-SCT antibody production were almost artificially and constitutively expressed in the cells, we also examined if endogenous glycosyltransferases faced the same problem of reduced glycosylation. By expressing antibody in FUT8 KO cells, the antibody was almost terminally glycosylated with at least one or more galactoses in the first collection (Fig. S6B, D0-3), but the proportion of galactosylated antibody decreased and even the antibody glycans without any galactoses became the dominant glycoforms in the last two days ((Fig. S6B, D4-5). The result showed that short glycosylation after protein overexpression can be observed on endogenous glycosylation products as well. This finding promoted us to dissect the impact of antibody production at different times on glycosylation. We purified antibody every day after plasmid transfection and analyzed the dynamic change of antibody glycans from multiple engineered cells. To our surprise, the antibody with full sialylation could be significantly detected by intact protein mass analysis at the first day, although it became unclear after that (Fig. 6). Overall, the most abundant sialylation level was observed at first day and then started to decrease day by day (Fig. 6 and S4A). Regarding galactosylation, the antibody glycans were capped by at least two galactoses (2∼4 galactoses in total) on the first day, but started to decline on the second day, as evidenced by the appearance of one galactose on antibody glycans (Fig. 6). These observations suggest that antibody glycosylation was almost complete at the beginning of overexpression and gradually decreased over time.

**Figure 5.**
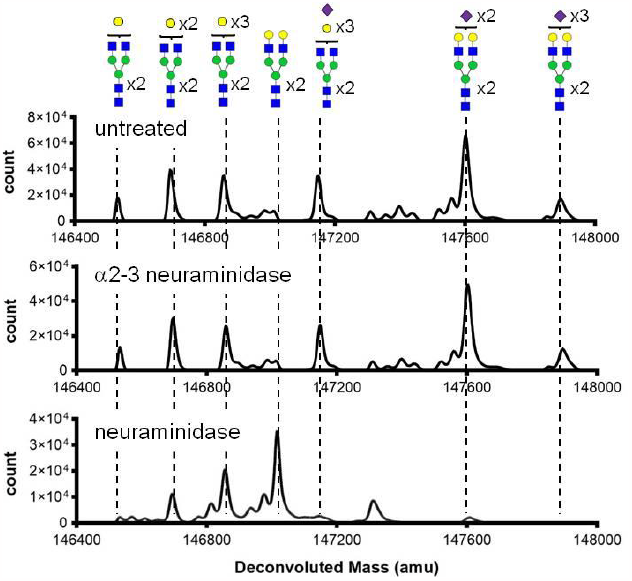
The sialylation on antibody is α2-6 linkage. The SCT enriched chMC81370 antibody was expressed by glycsylation pathway engineered cells. After incubation with indicated enzyme overnight, the antibody was purified by protein A and analyzed by intact protein mass analysis.

**Figure 6.**
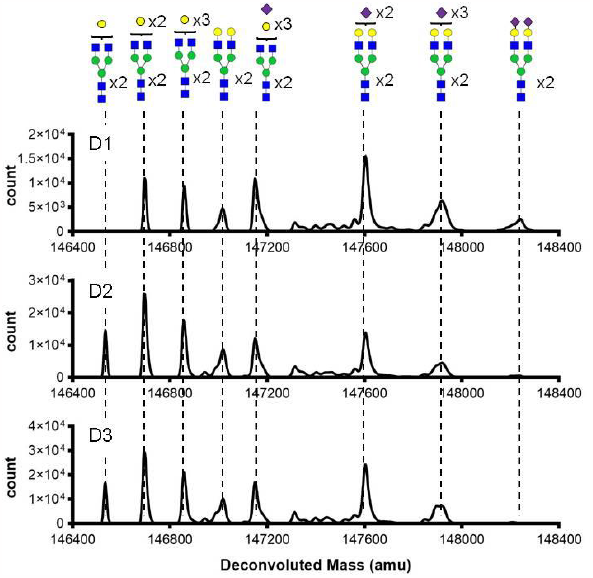
Antibody expression and time-dependent changes of glycans on antibody. The chMC81370 antibody expressed in glycosylation pathway engineered cells was collected every day and purified by protein A for intact protein mass analysis.

### Fc-SCT enriched antibody showed better binding to the Fcγ receptors than WT antibody

As we can obtain Fc-SCT enriched antibody from glycosylation pathway engineered cells, the next step was to compare the binding of the mixture of antibody glycoforms to Fc receptors with WT antibody^3^. FcγIIA and FcγIIB are usually expressed simultaneously by the antigen presenting cells, such as dendritic cells and macrophage, and these two receptors work together to modulate immune response^3^. FcγIIIA is critical for NK cells to mediate ADCC^3^. Therefore, the binding of antibody to these receptors was evaluated. We did not study the binding to FcγIA here because the glycoforms of complex type *N*-glycans on antibody would not affect the binding^23-24^. We observed the binding of Fc-SCT enriched antibodies to FcRs was increased in three tested Fc receptors (Fig. 7). Among these receptors, the increased binding of FcγIIA and FcγIIIA was consistent with previous studies^23-24^ (Fig. 7A and 7C). Interestingly, the WT antibody showed very weak binding to inhibitory receptor FcγIIB. Although the engineered *N*-glycans on AbHC improved binding to FcγIIB, the improvement was less significant than binding to FcγIIA (Fig. 7B). This observation was aligned with the fact that both FcγIIB and α2-6 sialylation were the key factors in intravenous Ig (IVIg) treatment-mediated inflammatory suppression^36-38^. Although the increased binding to the two opposing functional receptors, FcγIIA and FcγIIB, which are typically expressed on the antigen-presenting cells (APCs) simultaneously, appears to be conflicting, the eventual immune response was determined by the expression level and signaling strength of FcγIIA and FcγIIB on effector cells^3^. The Fc-SCT enriched antibody also exhibited stronger FcγIIIA binding ability than the antibody produced from wild type cells (Fig. 7C). This enhancement was mainly contributed by removal of core-fucose^11^ and addition of galactoses^23^.

**Figure 7.**
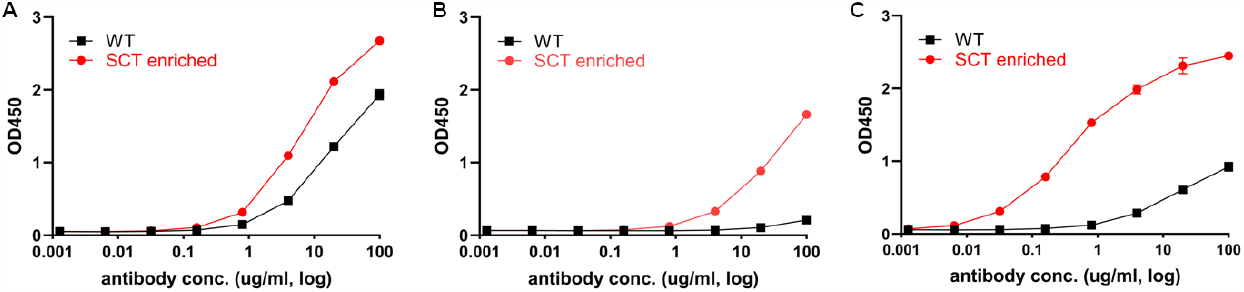
Improved receptor binding of SCT enriched antibody. The Fcγ receptor-coated plate (A, FcγIIA; B, FcγIIB; C, FcγIIIA) was incubated with WT antibody (black) or SCT enriched antibody (red) at indicated concentrations. The binding of antibody was measured by anti-human IgG Fc antibody conjugated with HRP. The binding of SCT enriched antibody to tested receptors was greatly improved compared with WT.

### Fc-SCT enriched antibody and homogenous Fc-SCT antibody are similar in functions

Since the binding of Fc-SCT enriched antibody to FcγIIIA was elevated, we compared the binding with the homogeneous Fc-SCT antibody. The result showed that Fc-SCT enriched antibody exhibited similar binding avidity to homogenous SCT antibody (Fig. 8A). Then we further evaluated if the Fc-SCT enriched antibody could evoke similar effect as homogeneous Fc-SCT antibody in ADCC reporter assay^39^. At the lowest antibody concentration used for induction, WT antibody did not induce any obvious cell activation, but the homogenous Fc-SCT and Fc-SCT enriched antibodies can trigger cell activation and rapidly reached a maximum at next concentration (Fig. 8B). Even at the higher concentration, the WT antibody can reach a maximum induction, but the activity was still much lower than that of homogenous Fc-SCT or Fc-SCT enriched antibody (Fig 8B). The overall response induced by homogenous Fc-SCT and Fc-SCT enriched antibodies were also very similar in receptor binding assay (Fig 8B). Thus, we concluded that the Fc-SCT enriched antibody derived from glycsylation pathway engineered cells could improve antibody effector functions in a similar manner as homogenous Fc-SCT antibody did.

**Figure 8.**
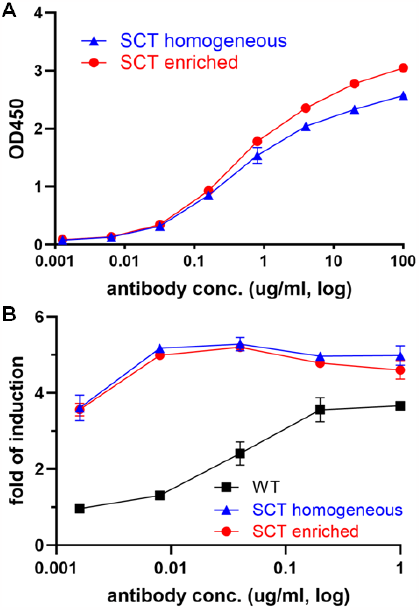
Functional comparison of homogenous SCT and SCT enriched antibody. (A) The FcγIIIA-coated plate was incubated with homogeneous SCT antibody (blue) or SCT enriched antibody (red) at indicated concentration. The binding of antibody was measured by anti-human IgG Fc antibody conjugated with HRP. The binding of SCT enriched antibody and homogenous SCT antibody to FcγIIIA was similar. (B) Indicated antibody, target cells, and effector cells were co-incubated for 6 hours followed by detection of luciferase activity. SCT enriched and homogenous antibody SCT showed similar effects on induction of ADCC reporter response and both elicited higher response than WT antibody.

In summary, we have simplified the process in different ways to generate functional Fc-SCT antibodies and evaluated the utility. First, we demonstrated that with endoglycosidase S2 or H expressed in the glycosyltranferase KO cells, the glycoengineered cells can produce antibodies with Fc-GlcNAc and the antibody made by this way can be used for *in vitro* enzymatic transglycosylation (Fig. 2-4). This strategy can be applied to engineer other *N*-glycsoylated proteins to produce GlcNAc-decorated glycoproteins as well (Fig. 5). Second, we knocked out the core-fucosyltransferase and constitutively expressed the glycosytransferases required for the enrichment of Fc-SCT glycan for cell-based production of glycoproteins. The Fc-SCT enriched antibodies generated by these cells (Fig. 6) exhibited similar FcR binding profiles to homogeneous Fc-SCT antibody and was able to activate similar effector functions such as ADCC through binding to FcγIIIA (Fig. 9). Thus, the cell-based production of Fc-GlcNAc and Fc-SCT enriched antibodies may have the potential advantage of simplifying the production of therapeutic antibodies with improved effector functions, especially ADCC, ADCP and vaccinal effect. Work in progress to evaluate the comparative activities of Fc-SCT enriched antibody glycoforms and homogeneous antibody in animals

## ASSOCIATED CONTENT

### Supporting Information

This supporting information is available free of charge via the Internet at http://pubs.acs.org, including general methods and materials, IPM analysis of antibody glycan, observation of survival rate, receptor binding of antibody.

## AUTHOR INFORMATION

### Author Contributions

H.-W. H. and C.-H. W. designed research; H.-W. H. performed research and analyzed data; V. S. and T.-H. T assisted trans-glycosylation assay; H.-W. H and C.-H. W wrote the paper.

### Notes

The authors declare no competing financial interest

## ACKNOWLEDGMENT

We would like to thank Han-Chung Wu for providing the plasmids of chMC-813-70 antibody, the mass core facility in the Scripps Research Institute for intact protein mass analysis, and the glycoscience core facility in Academia Sinica for technical supports of glycoform analysis. The Glycoscience Core Facility is funded by the Academia Sinica Core Facility and Innovative Instrument Project (AS-CFII-112-102). This work was supported by the National Institutes of Health (RO1AI130227) and the National Science Foundation (to CHW).

## Supplementary Information

### Methods and Material

#### Cell Culture

HEK 293T cells (ATCC) were cultured in the DMEM (Thermo Fisher) supplied with 10% FBS (Thermo Fisher) at 37°C with 5% CO_2_ in the cell incubator. EXPI293F and EXPI293F GNT1 KO cells (Thermo Fisher) were cultured in the EXPI293 expression medium (Thermo Fisher) at 37°C with 8% CO_2_ in the cell incubator.

#### Plasmid

The plasmids for CRISPR/Cas9 expression were constructed as kit manual (Thermo Fisher). Briefly, the validated sgRNA sequence targeting glycosyltransferase^1^ was synthesized (Integrated DNA Technologies) and cloned into the vector for Cas9 and sgRNA expression. The synthetic codon-optimized gene insert (Integrated DNA Technologies), flanked by homologous arm of target gene at sgRNA target site, was cloned into empty vector as the donor plasmid.

#### Transfection

Transfection of 293T cells and EXPI293F cells were mediated by TransIT-293 (Mirus Bio) or expi293fectamine (Thermo Fisher) following the reagent manual.

#### Antibody production

The plasmids for antibody chMC81370 were transfected into cells followed by incubation. At the time of harvest, the culture medium was collected and then subjected to protein A sepharose beads (GE HealthCare) column to purify the antibody.

#### Protein gel analysis

The protein was heated at 95 °C for 5 mins in the LDS sample buffer supplied with 2-Mercaptoethanol and then subjected to gel electrophoresis with 12% SDS-PAGE. The protein on gel was stained by coomassie blue buffer (ApexBio).

#### Enzymatic transglycosylation assay

The antibody was incubated with SCT glycan-oxazoline and Endo S2 mutant in Tris buffer at 37 °C for the time indicated^2^ before harvest for purification or gel electrophoresis.

#### Intact protein mass analysis

The samples were diluted with LCMS grade water at a concentration between 5-10uM and analyzed by 6230 TOF LC/MS with a Dual AJS ESI ion source (Agilent Technologies) with PLRP-S 1000Å 5µm column (Agilent Technologies). Solvent A was 0.1% Formic Acid in H2O and Solvent B was 0.1% Formic Acid in ACN.

#### Glycosidase treatment

The proteins were incubated with alpha-2-3 neuraminidase, neuraminidase, or PNGase F (New England Biolabs) in the supplied buffer at 37 °C overnight followed by purification or gel electrophoresis.

#### ELISA binding assay

The recombinant soluble FcγRIIIA (R&D) was coated with 50 ng/well in bicarbonate/carbonate coating buffer (50 mM, pH. 10) at 4°C overnight before blocking the well by 5% BSA in TPBS (0.05% Tween 20 in PBS) at 4°C overnight. The antibody was added into wells with final concentration started at 100 μg/ml with 5x series dilution and incubated at room temperature for 1 hour followed by incubation with HRP-conjugated goat anti-human IgG antibody (Jackson ImmunoResearch) for another 1 hour at room temperature. Finally, the TMB substrate (Bethyl Laboratories) was added to react with HRP at RT prior to stopping the reaction by H_2_SO_4_. The absorbance at 450 nm was detected by SpectraMax M5 spectrum reader (Molecular Device). The wells were washed by TPBS 3-5 times between each step.

#### ADCC reporter assay

ADCC reporter assay (Promega) was performed as kit manual. Briefly, the SSEA4 expressing target cells SK-OV3^3^ was seeded into 96-well plate followed by addition of antibody with final concentration started at 1 g/ml with 5x series dilution. Then the FcγRIIIA expressing effector cells (effector: target cell ratio, 6:1) was added and incubated with target cells in the incubator at 37 °C for 6 hrs. The plate was placed at room temperature for 15 min prior to adding the luciferase substrate. After 5 min incubation, the luminescence was measured by SpectraMax M5 spectrum reader. The induction fold was calculated by RLU (induced–background) /RLU (no antibody control–background).

#### Glycoform analysis

The FASP (filter-aided sample preparation) method^4^ was used for in-solution protease digestion. Briefly, 5 μg of protein samples were loaded onto filter units (Microcon YM-30) and mixed with 100 μl of 8 M urea in 0.1 M Tris-HCl pH 8.5 (UB) each for protein denaturation. The filter units were centrifugated with 14,000 x g for ∼30 min until the solution was removed, followed by the addition of 100 μl of 25 mM DTT in UB each for protein reduction, and incubated at room temperature for 10 min. After centrifugation with 14,000 x g for ∼30 min to remove the DTT solution, 100 μl of 50 mM iodoacetamide in UB was added to each sample for alkylation, and the filter units were incubated in the dark for 10 min, followed by centrifugation to remove iodoacetamide solution. To each filter unit, 100 μl of UB was added again. After another centrifugation to remove the solution, 100 μl of 50 mM ammonium bicarbonate in water (ABC) was added to change the buffer system to ABC for in-solution protease digestion. After centrifugation to remove the solution, 40 μl of ABC with 0.1 μg Trypsin/Lys-C Mix (Promega, sequencing grade) was added to each sample and incubated in a wet chamber at 37°C overnight. The digested peptides in samples were spun to collection tubes, and an additional rinse with 40 μl ABC for each sample was performed. The filter units were centrifuged at 14,000 x g for 10 min to collect the flow-through, which contains digested peptides. Samples were acidified with FA to a final concentration of 0.1%. The peptide samples were dried in a SpeedVac evaporator, and stored at -20°C until analysis with LC-MS/MS.

Samples were analyzed using the LC-ESI-MS method on an Orbitrap Fusion mass spectrometer (Thermo Fisher Scientific, San Jose, CA) equipped with an EASY-nLC 1200 system (Thermo, San Jose, CA, US) and an EASY-spray source (Thermo, San Jose, CA, US). A 5 μl digestion solution was injected at a flow rate of 1 μl/min onto an easy column (C18, 0.075 mm x 150 mm, ID 3 μm; Thermo Scientific). Chromatographic separation utilized 0.1% formic acid in water as mobile phase A and 0.1% formic acid in 80% acetonitrile as mobile phase B, operated at a flow rate of 300 nl/min. The gradient employed was from 5% buffer B at 2 min to 60% buffer B at 55 min. For full-scan MS, the conditions included a mass range of m/z 375-1800 (AGC target 5E5) with a lock mass, resolution set at 60,000 at m/z 200, and a maximum injection time of 50 ms. MS/MS was conducted in top speed mode with 3 s cycles using both CID and HCD, with a dynamic exclusion duration of 60 s and a 10 ppm tolerance around the selected precursor and its isotopes. The electrospray voltage was maintained at 1.8 kV, and the capillary temperature was set to 275°C.

Byonic software version 4.2.10 (Protein Metric) was used for the identification of summary formulas of glycans associated with glycopeptides. The glycan database contained 132 entries, and the parameters used were: precursor mass tolerance of 10 ppm, fragment mass tolerance of 0.5 Da, maximum missed cleavages set to 5, and cysteine carbamidomethylation and methionine oxidation considered.”

**Figure S1.**
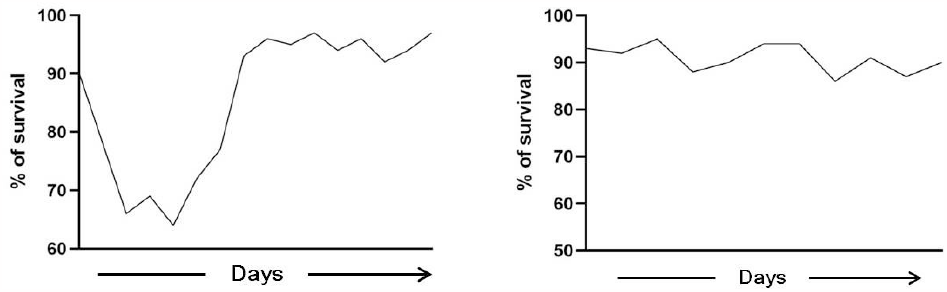
The survival curve of the GNT1-KO EXPI293F cells after CRISPR/Cas9 and Endo H (left) or Endo S2 (right) donor plasmid cotransfection.

**Figure S2.**
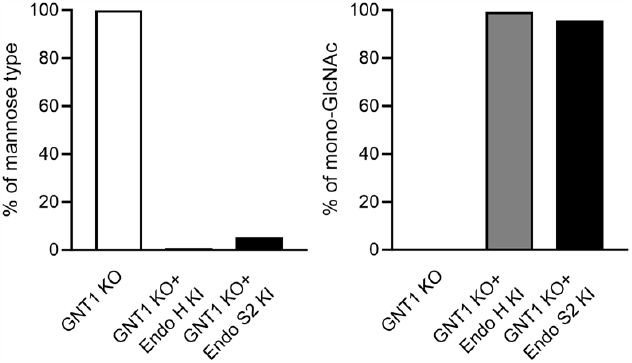
The glycoform analysis of chMC81370 antibody expressed by GNT1 KO and engineered GNT1 KO EXPI293 cells.

**Figure S3.**
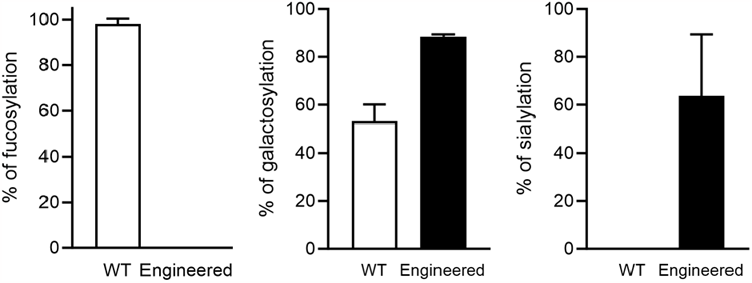
The glycoform analysis of chMC81370 antibody expressed by WT or engineered 293T cells.

**Figure S4.**
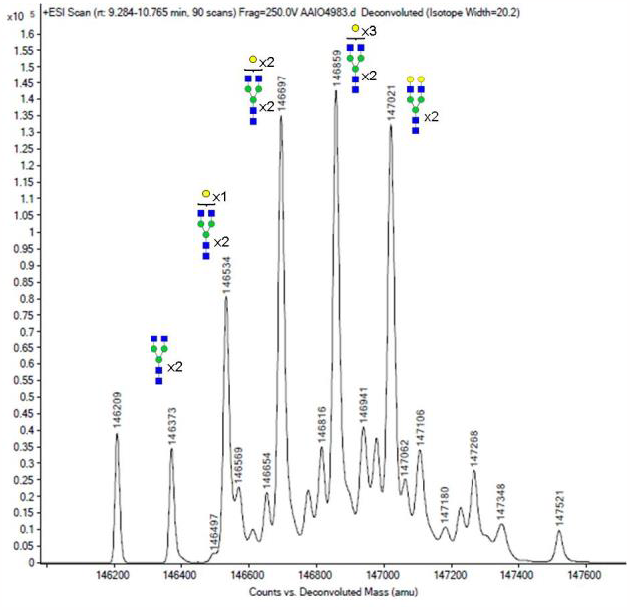
The IPM analysis of chMC81370 antibody expressed by FUT 8KO/B4GalT1 KI 293T cells

**Figure S5.**
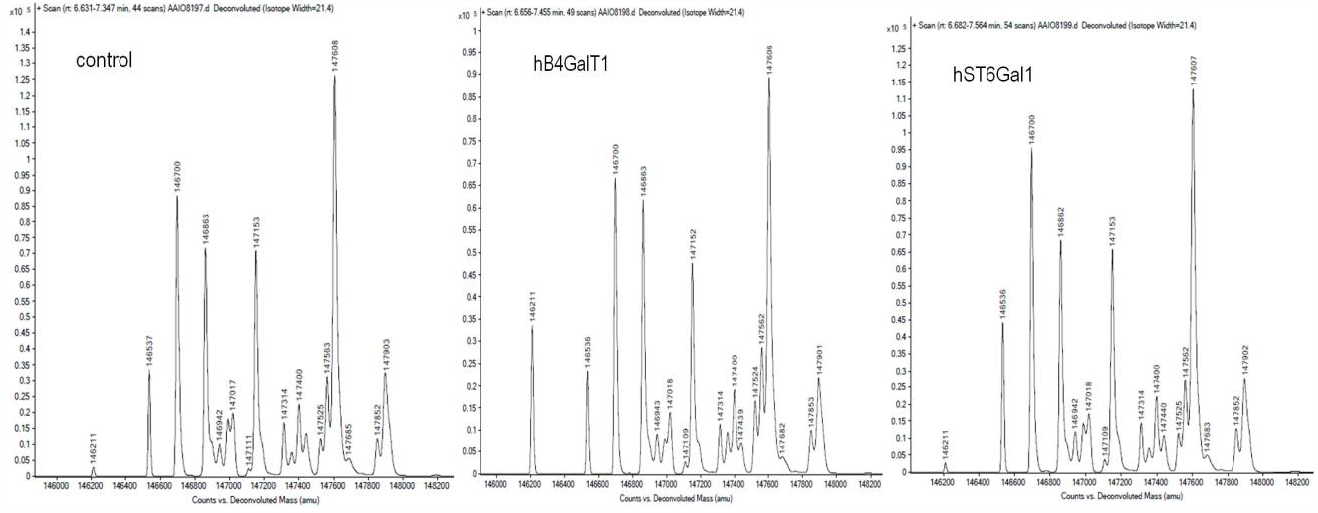
The IPM analysis of chMC81370 antibody expressed by untreated, hB4Gal1 transiently overexpressed, and hST6GalT1 transiently overexpressed, engineered 293T cells.

**Figure S6.**
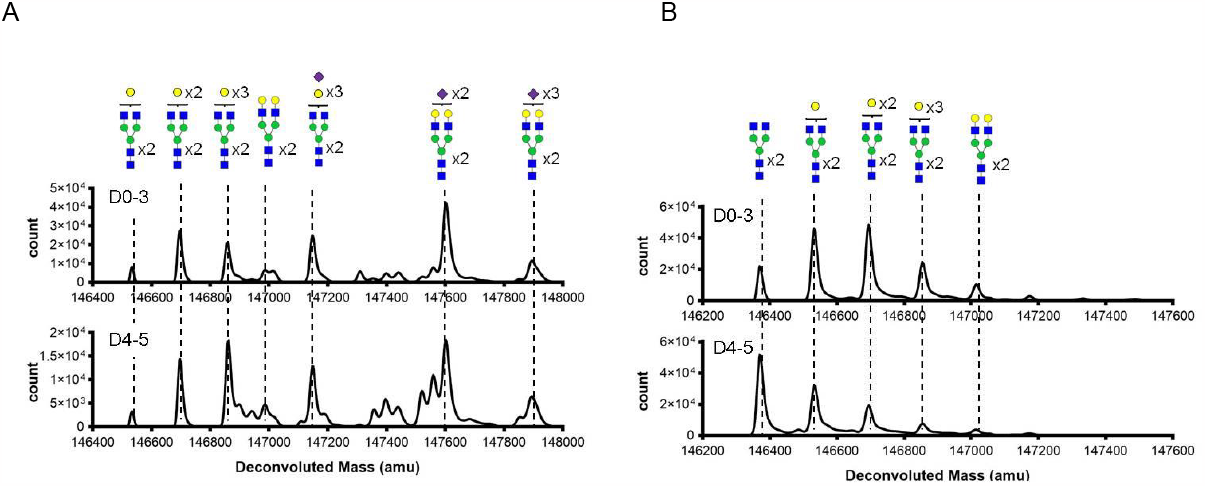
The IPM analysis of chMC81370 antibody expressed by multiple engineered (A) and FUT8 KO (B) cells at different collection time point.

